# Structural basis of ZP2-targeted immunocontraception

**DOI:** 10.1101/2025.03.03.641048

**Authors:** Elisa Dioguardi, Alena Stsiapanava, Eileen Fahrenkamp, Ling Han, Daniele de Sanctis, José Inzunza, Luca Jovine

**Author notes:** To whom correspondence may be addressed: Luca Jovine. Address: Department of Medicine, Karolinska Institutet, 14183 Huddinge, Sweden. E.D. and A.S. contributed equally to this work. **Competing Interest Statement:** The authors declare no competing interest.

## Abstract

Monoclonal antibody IE-3 prevents mouse fertilization by binding ZP2, a major component of the oocyte-specific zona pellucida (ZP). We show that an IE3-derived single-chain variable fragment (scFV) is sufficient for blocking fertilization in vitro and determine the structural basis of IE-3/ZP2 recognition. The high-affinity of this interaction depends on induced fit of the epitope, offering insights for non-hormonal immunocontraceptive design without off-target effects.

## Introduction

Due to significant technological advances, there is a renewed interest in developing contraceptive antibodies that prevent fertilization by targeting gamete interaction proteins (1). As the first interface between egg and sperm (2), the ZP has long been explored as a target for immunocontraception (3), with notable proof-of-principle applications to wildlife control (4, 5). Among the monoclonal antibodies raised against ZP proteins, of particular interest is IE-3, which targets the N-terminal ZP-N1 domain of mouse ZP2 (mZP2-N1) (6, 7). This domain was suggested to directly mediate sperm binding to the ZP (7) and implicated in the block to polyspermy by regulating the post-fertilization cleavage of ZP2 (8). IE-3 binds to the ZP in vitro and in vivo (9–11) and has a full contraceptive effect in mice, with a reversible block of animal fertility (9) and no interference with normal follicular maturation (10). These properties establish IE-3 as a solid foundation for developing non-hormonal contraceptives. However, how the antibody exactly recognizes ZP2 at the molecular level is unknown and, more generally, there is no structural information on any antibody/fertilization protein complex.

## Results and Discussion

To study how IE-3 recognizes its epitope on ZP2, we sequenced the antibody gene from its hybridoma line and engineered a His-tagged scFV that was expressed in mammalian cells and purified by immobilized metal-ion affinity chromatography (IMAC). Size-exclusion chromatography (SEC) showed that scFV forms a stable complex with mZP2-N1 in solution (Fig. 1*A*). In vitro fertilization (IVF) experiments showed that oocytes pre-incubated with scFV remained unfertilized, whereas oocytes exposed to buffer or human growth hormone (hGH; a control His-tagged glycoprotein of comparable molecular weight) developed into embryos (Fig. 1*B, C*). Notably, scFV did not affect sperm motility but blocked fertilization by both reducing the attachment of sperm (Fig. 1*D*) and hindering ZP penetration, as suggested by the lack of spermatozoa within the ZP or in the perivitelline space (Fig. 1*C*).

**Fig. 1.**
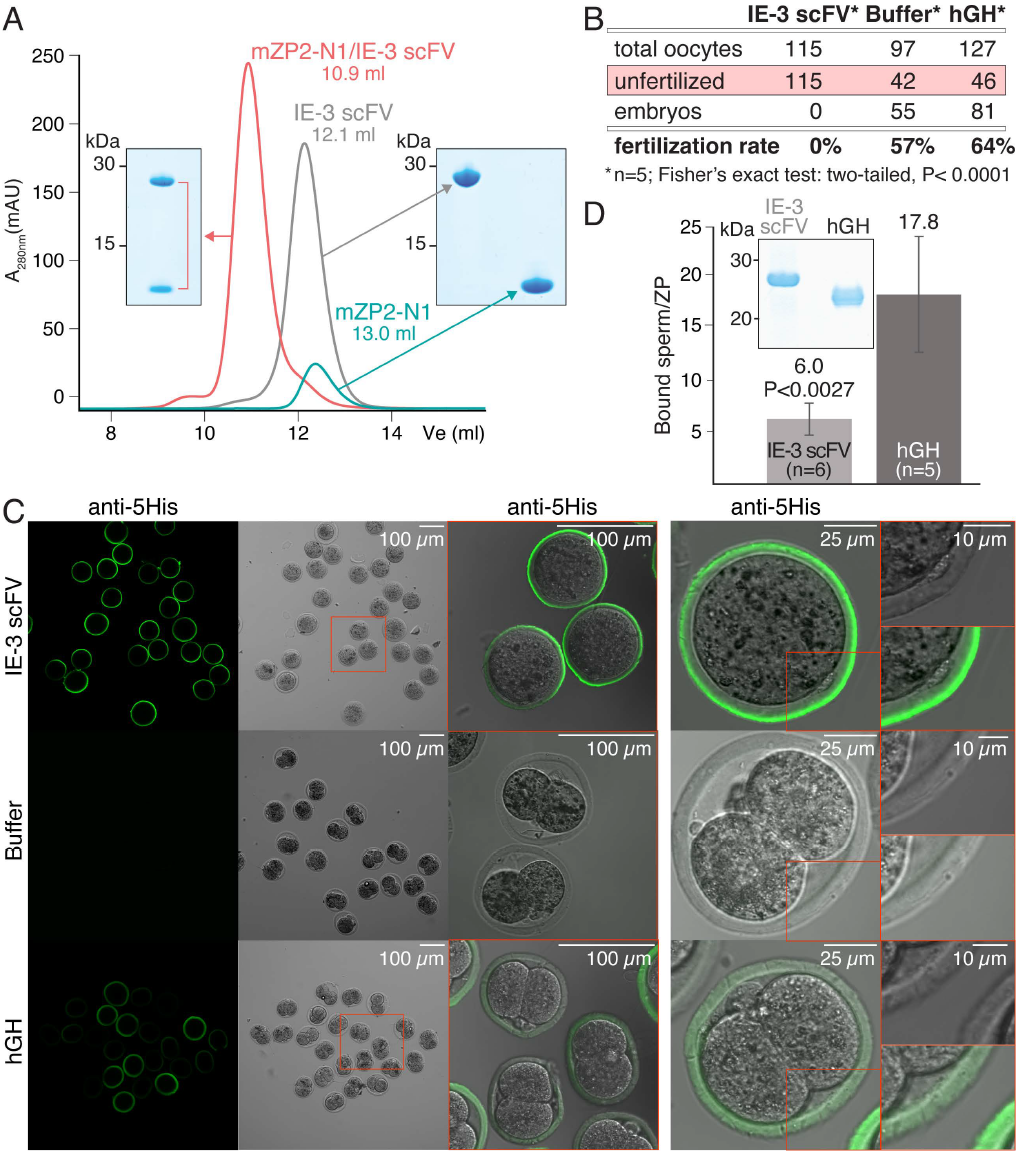
IE-3 scFV binds recombinant mZP2-N1 and blocks fertilization in vitro. *(A)* SEC analysis of purified mZP2-N1, scFV or their complex. V_e_, elution volume. *(B)* Fertilization rates of oocytes incubated with scFV, buffer or hGH. *(C)* Anti-5His immunofluorescent staining of representative scFV-, buffer- or hGH-treated oocytes used for the IVF experiments summarized in panel *B*. hGH-treated oocytes show a weak fluorescent signal due non-specific protein binding to the ZP. Only oocytes incubated with buffer or hGH developed into 2-cell embryos. *(D)* Number of ZP-bound sperm after oocyte treatment with purified scFV or hGH (inset).

To structurally investigate antibody-protein recognition, we co-expressed the unfused variable heavy and light chains of IE-3 (V_H_, V_L_) together with His-tagged mZP2-N1. This enabled efficient secretion and purification of the trimeric complex, which produced two crystal forms whose structures were determined to 1.5 Å and 2.1 Å resolution, respectively. The corresponding models are essentially identical (root-mean-square deviation (RMSD) 0.28 Å) and their superposition onto the structure of the full N-terminal region of human ZP2 (hZP2 NTR) (8) shows that, consistent with the ability of IE-3 to recognize ZP2 in native conditions (12), the V_H_V_L_ heterodimer binds to the surface of ZP2-N1 opposite to its junction with the rest of the molecule (Fig. 2*A*). The three complementarity-determining regions of V_H_ (CDR-H1/H2/H3), together with CDRs L1 and L3 of V_L_, generate a hydrophobic pocket where the fg loop of mZP2-N1 inserts. This dominates the interaction with IE-3, which includes only a few other contacts between the cd loop of mZP2-N1 and CDR-L1 (Fig. *2A*).

**Fig. 2.**
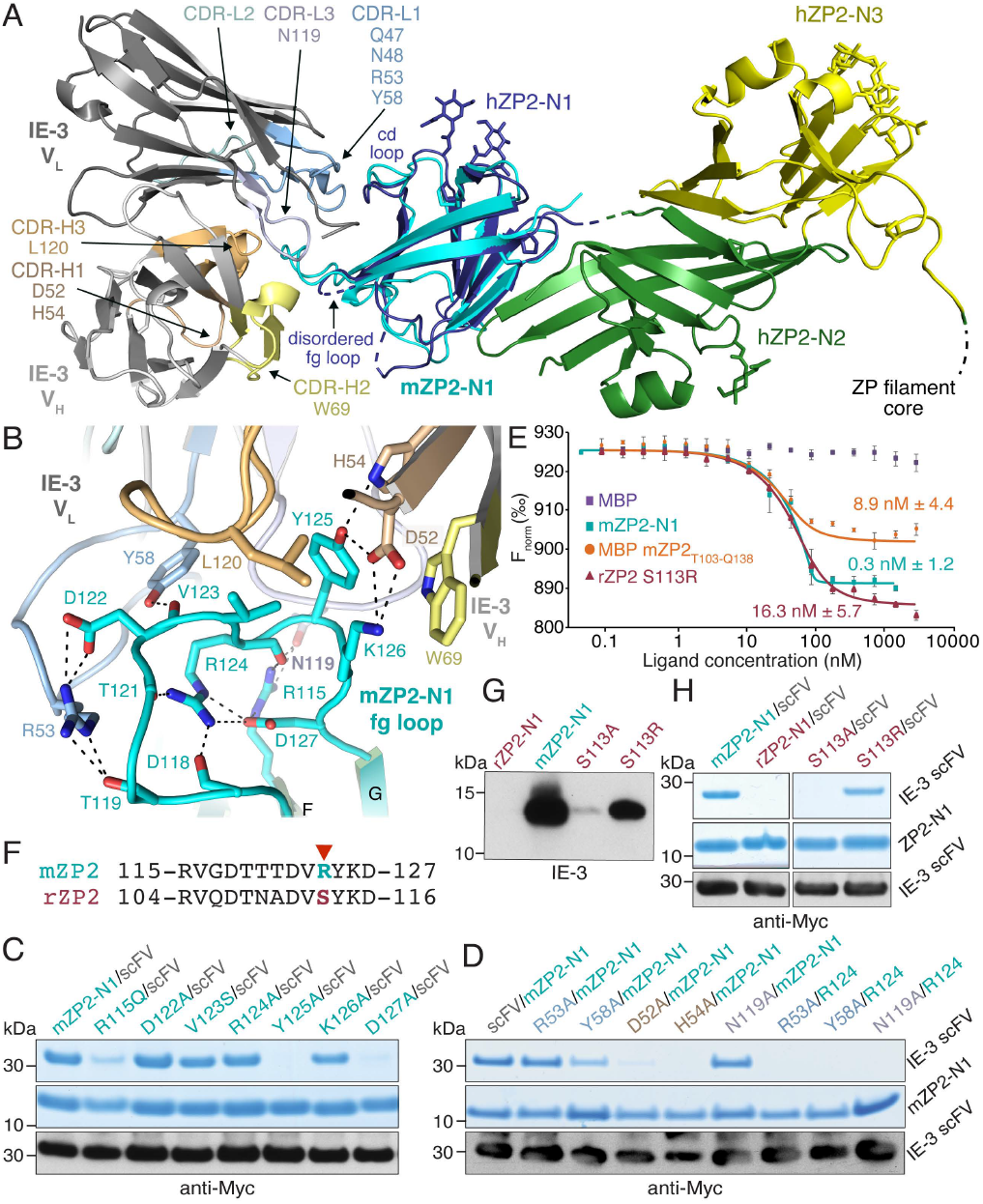
Structural analysis of mZP2-N1 recognition by IE-3. *(A)* Superposition of the mZP2-N1/V_H_V_L_ complex structure onto hZP2-N1N2N3 (PDB ID: 8RKE) with RMSD of 1.25 Å across 78 aligned residues. Selected CDR residues at the interface are indicated. *(B)* Detail of mZP2-N1 fg loop/V_H_V_L_ CDR interactions. Dashed lines indicate hydrogen bonds; water molecules are omitted for clarity. *(C-D)* Coomassie-stained SDS-PAGE analysis of pull-down experiments using mZP2-N1 mutants *(C)* or scFV mutants and mZP2-N1/scFV double mutants *(D)*. Immunoblots display the respective scFV input levels (100 µl). *(E)* Microscale thermophoresis (MST) analysis of the interaction between the fragment antigen-binding (Fab) region of IE-3 and mZP2/rZP2 constructs. MBP was used as a negative control. Kd values are reported (error bars = s.d.; n = 3). *(F)* Sequence alignment of rodent ZP2-N1 fg loops. A red arrow indicates mZP2 R124 and rZP2 S113. *(G)* Whereas wild-type rZP2-N1 does not bind IE-3, the S113R mutant shows significant binding. *(H)* Pull-downs of rZP2-N1 mutants show that S113R forms a stable complex with IE-3 scFV.

Notably, the long fg loop of ZP2 is largely disordered in the structures of both unbound mZP2-N1 (13) and hZP2 NTR (8) and becomes fixed upon interaction with the antibody (Fig. *2A, B*). Moreover, the IE-3-interacting region of the fg loop includes residues _123_VRYK_126_, which match the sequence VxYK that was previously identified as a minimal consensus epitope for IE-3 by screening a random peptide library (6). In agreement with this study, the structure of the complex and binding assays of point mutants show that hydrogen bonding of mZP2-N1 Y125 to CDR-H1 D52 and H54 is crucial for epitope recognition. On the other hand, mutation of fg loop K126, which makes a cation-π interaction with CDR-H2 W69 and also hydrogen bonds to CDR-H1 D52, does not affect complex formation (Fig. 2*B-D*). Notably, the structure reveals that F-β-strand R115 and fg loop D122, two residues located outside the VxYK motif, also contribute to mZP2-N1 recognition by contacting CDR-L3 N119 and CDR-L1 R53, respectively. Most importantly, the mZP2-N1/IE-3 interaction indirectly depends on D127, which does not contact the antibody but orients R115 as well as influences the overall conformation of the fg loop by stabilizing a set of interactions centered around R124 (Fig. 2*B*). Accordingly, complex formation is significantly reduced by either individually mutating mZP2-N1 R115 or D127, or combining a mZP2-N R124A substitution with a mutation of V_L_ R53, Y58 or N119 (with each of these mutations not being sufficient to disrupt binding by itself) (Fig. 2*C, D*). Consistent with an important role of the 3D shape of the fg loop, IE-3 binds mZP2-N1 with 30-fold higher affinity than the mZP2_T103-Q138_ linear epitope fused to maltose-binding protein (MBP) (Fig. 2*E*). Interestingly, IE-3 does not bind to the ZP2-N1 domain of rat ZP2 (rZP2), which contains all the aforementioned key residues except for a Ser at position 113, corresponding to mZP2-N1 R124 in the VxYK motif. However, in agreement with the above observations, mutation of S113 to Arg (but not Ala) allows rZP2-N1 to also be recognized by IE-3 scFV, albeit with reduced affinity (Fig. 2*E-H*).

Our results have three main implications for the development of ZP2-targeted contraceptives. First, the finding that, despite its relatively small size, an IE-3 derived scFv is sufficient to block fertilization in vitro identifies a framework that will intrinsically avoid potential Fc-associated side effects. Second, the high-resolution structural information on mZP2-N1/V_H_V_L_, reveals that the 3D conformation of the epitope contributes significantly to its high-affinity recognition by the antibody. Consistent with the fact that IE-3 was obtained by immunization using native ZPs (12), this underlines the importance to use properly folded ZP2 material as antigen, as opposed to using linear ZP2-derived peptide sequences (6) or heat-solubilized pig ZP material (4). Third, the structure shows that IE3 CDR-L1 makes auxiliary contacts with the cd loop of mZP2-N1. Although our experiments exclude a requirement for these interactions, rational design of CDR-L1 variants could be used to further increase the affinity for ZP2.

## Materials and Methods

Proteins were expressed in human embryonic kidney 293 cells and purified by IMAC and SEC. IVF assays were performed using oocytes from juvenile C57BL6/J female mice and cauda epididymis sperm from mature male mice. Diffraction data was collected at beamline ID29, European Synchrotron Radiation Facility (France). Binding affinities were determined by MST. A full description of the methodology is detailed in *SI Appendix, Materials and Methods*.

## Data, Materials, and Software Availability

X-ray diffraction and structural data have been deposited in Protein Data Bank (9H4R (14), 9H4S (15)).

## Acknowledgments

This research was supported by Swedish Research Council grants 2012-05093, 2016-03999, 2020-04936, European Research Council grant 260759, Knut and Alice Wallenberg Foundation grant 2018.0042 (L.J.); European Molecular Biology Organization long-term fellowship 143-2017 and Deutsche Forschungsgemeinschaft research fellowship DI 2224:2-1 (E.F.).

## Supplementary Information

## Supplementary Materials and Methods

### DNA constructs

The details of the mZP2-N1 construct used in this study have been described previously (1). For producing additional ZP2 proteins used for binding assays, a construct encoding C-terminally 6His-tagged rZP2-N1 (rZP2_M1-Q127_ with N-glycosylation site mutation N72S) was synthesized (Life Technologies/Thermo Fisher Scientific) and cloned into mammalian expression vector pHLsec3 (2); additionally, a cDNA fragment encoding mZP2 T103-Q138, essentially corresponding to the protein’s B cell epitope (3), was subcloned in frame with the MBP ORF of pLJMBP6 (4).

The cDNAs for the IE-3 heavy and light chains (HC/LC) were amplified by RT-PCR from the hybridoma cell line CRL-2463 (5) (ATCC) and sequenced (Genscript); the corresponding sequences have been deposited with GenBank under accession numbers MH212328 and MH212329, respectively. Note that the V_L_ sequence used in this study differs from that included in a previously described synthetic construct (6) at 5 positions (T71, H75, W76, Q81 and S82, corresponding to GenBank entry ALJ49777 K579, S583, L584, E589 and T590, respectively). Although none of these residues is in contact with ZP2, W76 stabilizes the V_H_/V_L_ interface by interacting with V_H_ Y122, at the periphery of the hydrophobic pocket that binds the mZP2-N1 fg loop.

For IE-3 Fab expression, cDNAs encoding HC_M1-R239_ followed by a 6His-tag and LC_M1-C240_ were cloned into pHLsec3; for IE3 V_H_V_L_ expression, HC_Q20-S139_ and LC_D21-R134_ were cloned in frame with the Crypα signal peptide of pHLsec3. To generate the construct encoding IE-3 scFV, V_L_ and V_H_ sequences were connected by a 16-residue (GGGS)_4_ linker and followed by 6His or Myc tags. For expression of the C-terminally His-tagged hGH control protein, the corresponding gene insert was amplified by PCR from pSGHV0 (7) and cloned into pHLsec3.

All mutations were introduced by overlapping PCR. Oligonucleotides were ordered from Sigma-Aldrich and all constructs were verified by DNA sequencing (Eurofins Genomics).

### Protein expression

Proteins were transiently expressed in human embryonic kidney (HEK) 293 cells using 25 kDa branched polyethylenimine (2). For co-expression experiments of mZP2-N1 and IE-3 V_H_V_L_ in HEK293T cells, 3 ng/μl mZP2-N1 DNA were mixed with 3.5 ng/μl DNA V_H_ and 3.5 ng/μl DNA V_L_; all other co-transfections were performed using a 1:1 DNA ratio.

MBP-mZP2_T103-Q138_ and control MBP were expressed in *E. coli* BL21 pLysS (DE3) (Promega) at 21ºC. Protein expression was induced for 16–18 h with 0.1 mM IPTG at an optical density (OD_550_) of 0.5–1. Cells from 1 L culture, suspended in 10 mL 50 mM Tris–HCl pH 7.5, 50 mM NaCl, 1 mM MgCl_2_, 0.2 mg mL^-1^ lysozyme, 25 U mL^-1^ Benzonase (Sigma-Aldrich) and cOmplete mini EDTA-free protease inhibitors (Roche), were disrupted using three freeze-thaw cycles. Bacterial debris was removed by centrifugation at 18,000 x g for 30 min.

### Protein purification

72 h after transfection, the conditioned media from mammalian cells was harvested, 0.22 μm-filtered (Sarstedt) and adjusted to 20 mM Na-HEPES pH 7.8, 500 mM NaCl, 5-10 mM imidazole (IMAC buffer). 10 mL pre-equilibrated nickel agarose slurry (Ni-NTA; QIAGEN) was added per liter of medium and incubated for either 1 h at room temperature (RT). After washing the beads with 100 column volumes IMAC buffer, proteins were batch-eluted with 5 column volumes elution buffer (20 mM Na-HEPES pH 7.8, 150 mM NaCl, 500 mM imidazole). Proteins were concentrated with appropriate centrifugal filtration devices (Amicon) and further purified by SEC at 4ºC using an ÄKTA_FPLC_ chromatography system (GE Healthcare). mZP2-N1/IE-3 V_H_V_L_ complex was injected into a Superdex 75 26/60 column (GE Healthcare) pre-equilibrated with 10 mM Tris-HCl pH 7.2, 50 mM NaCl. Peak fractions were pooled and the protein complex was concentrated to 27 mg mL^-1^ for crystallization trials.

Batch IMAC of MBP-mZP2_T103-Q138_ and MBP were carried out for 1 h at RT in 50 mM Tris-HCl pH 7.5, 1 M NaCl, 20 mM imidazole; proteins were eluted with 50 mM Tris-HCl pH 7.5, 150 mM NaCl, 500 mM Imidazole and further polished by SEC using a Superdex 200 26/600 column (GE Healthcare) pre-equilibrated with 10 mM Na-HEPES pH 7.5, 150 mM NaCl.

### Protein analysis

For immunoblotting, proteins separated by SDS-PAGE were transferred to a nitrocellulose membrane (GE Healthcare Life Sciences) and probed with the following primary antibodies: anti-mZP2 IE-3 monoclonal (1:1,000); anti-5His monoclonal (1:1,000; QIAGEN); anti-Myc monoclonal (1:1,000; Sigma-Aldrich clone 9E10). Secondary antibodies were goat anti-rat (1:10,000; Thermo Fisher Scientific); horseradish peroxidase-conjugated goat anti-mouse (1:10,000) (Life Technologies/Thermo Fisher Scientific). Chemiluminescence detection was performed with Western Lightning ECL Plus (Perkin Elmer).

### Pull-down analysis of protein-protein interaction

2 mL conditioned medium containing 6His-tagged mZP2-N1 were harvested three days post-transfection and centrifuged for 5 min at 500 x g. 20 mM Na-HEPES pH 7.8, 150 mM NaCl and 5 mM imidazole were added and samples were incubated with 50 μL Ni-NTA beads for 1 h at RT. Beads were collected by centrifugation at 100 x g and washed 3 times with 1 mL 20 mM Na-HEPES pH 7.8, 150 mM NaCl, 10 mM imidazole.

Bound material was eluted with 100 μL 20 mM Na-HEPES pH 7.8, 150 mM NaCl, 500 mM imidazole and 20 µL were analyzed by immunoblot or Coomassie blue staining.

### Binding affinity determination by microscale thermophoresis

MST analysis was performed using a NanoTemper Monolith NT.115 instrument (NanoTemper Technologies). Because of severe bleaching of IE-3 scFV, recombinantly expressed IE-3 Fab was used and labeled with the Blue-NHS labeling kit (NanoTemper Technologies GmbH), according to the manufacturer’s instructions and using a labeling buffer containing 10 mM Na-HEPES pH 7.5, 150 mM NaCl, 0.01% (v/v) Tween 20.

Varying concentration of mZP2-N1, rZP2-N1 S113R and MBP-mZP2_T103-Q138_ in labeling buffer were titrated against labeled IE-3 Fab (8 nM) in 20 mM Na-HEPES pH 7.8, 200 mM NaCl, 0.01% (v/v) Tween 20. Unfused MBP was used as negative control. Samples were loaded into Premium Coated Capillaries (NanoTemper Technologies) and measurements were performed using 20% MST power and 85% LED power. Laser-on and -off times were 30 s and 5 s, respectively. Each of the graphs superimposed in Fig. 2E displays merged data from three independent experiments. Datasets were processed with the MO.Affinity Analysis software (NanoTemper Technologies) using the signal from the thermophoresis T jump.

### Protein crystallization

Crystallization experiments were performed by hanging drop vapor diffusion at 20ºC using a mosquito crystallization robot (TTP Labtech) using a Nunc 96 multiwell plate (Thermo Scientific) with a ratio protein:reservoir 1:1 (drop size 200 nL). The mZP2-N1/IE-3 V_H_V_L_ complex crystallized in 24% (v/v) PEG 3350, 0.2 M sodium tartrate, 0.1 M Na-HEPES pH 6.5 (crystal form I; *P*2_1_2_1_2_1_) as well as 2.0 M lithium sulfate, 0.2 M ammonium sulfate, 0.1 M tri-sodium citrate pH 5.6 (crystal form II; *P*4_3_).

### X-ray diffraction data collection

Crystals were cryoprotected using mother liquor solutions supplemented with 15% (v/v) PEG 200 (crystal form I) or 10% (v/v) PEG 200 (crystal form II).

All datasets were collected from single crystals at 100 K. The initial orthorhombic dataset of mZP2-N1/IE-3 V_H_V_L_ was collected in house using a Compact HomeLab system (Rigaku) equipped with a PILATUS 200K detector (DECTRIS), while final datasets for both crystal forms were collected at a wavelength of 0.9763 Å at European Synchrotron Radiation Facility (ESRF) beamline ID29 (8), using a PILATUS 6M-F detector (DECTRIS). For crystal form I, data collection statistics generated using phenix.table_one (9) were: completeness 93.4 (83.8) %, multiplicity 3.3 (3.4), mean I/sigma(I) 20.2 (1.1), R_merge_ 0.025 (1.166), R_meas_ 0.030 (1.375), R_pim_ 0.016 (0.720), CC_1/2_ 1.00 (0.54), CC* 1.00 (0.84) (with the values in parentheses being for the highest resolution shell, 1.58-1.53 Å). For crystal form II, statistics were: completeness 97.3 (92.5) %, multiplicity 6.8 (7.1), mean I/sigma(I) 26.1 (1.5), R_merge_ 0.052 (1.369), R_meas_ 0.056 (1.478), R_pim_ 0.021 (0.553), CC_1/2_ 1 (0.53), CC* 1.00 (0.83) (with the values in parentheses being for the highest resolution shell, 2.09 - 2.02 Å).

### Data processing and structure determination

X-ray diffraction datasets were processed using XDS (10). A 2.30 Å resolution dataset collected in house was used to determine the orthorhombic structure of the mZP2-N1/IE-3 V_H_V_L_ complex by molecular replacement (MR) with Phaser (11), using as independent search models the structure of unbound mZP2-N1 (PDB ID 5II6, chain A residues P43-Q138) (1) and those of the V_H_ domain of antiporphyrin antibody 13g10 (PDB ID 4AMK, chain H residues Q3-S118) (12) and the V_L_ domain of chimeric antibody X836 (PDB ID 3MBX, chain L residues D1-R114) (13), both of which were pre-processed with Sculptor (14). An initial model, consisting of two copies of the complex related by non-crystallographic symmetry (NCS) (corresponding to mZP2-N1, V_H_, V_L_ chains A, H, L and B, X, Y, respectively), was autotraced with PHENIX AutoBuild (15). After manual rebuilding in Coot (16), the model was refined to R/R_free_ 0.206/0.233 against the 1.53 Å resolution synchrotron dataset using phenix.refine (17) (with torsion-based NCS restraints, as well as twinning operator k, h, -l and twinning fraction 0.3). Protein geometry and all-atom contacts were validated with MolProbity (18): Ramachandran favored/allowed/outliers: 98.2/1.8/0.0 %; Rama distribution Z-score: 0.53 ± 0.31; favored/poor rotamers: 96.5/0.0 %; MolProbity score: 1.12; clashscore 3.28.

The tetragonal structure of mZP2-N1/IE-3 V_H_V_L_, whose asymmetric unit also includes two complexes, was solved by MR using a partially refined model of the *P*2_1_2_1_2_1_ structure. The solution was rebuilt, refined to R/R_free_ 0.189/0.203 at 2.02 Å resolution, and validated as above (Ramachandran favored/allowed/outliers: 98.0/2.0/0.0 %; Rama distribution Z-score: -0.08 ± 0.31; favored/poor rotamers:rotamer outliers: 98.1/0.2 %; MolProbity score: 1.00; clashscore 2.24). The average B-factor of V_L_ (chain L) in the orthorhombic structure of the complex is significantly higher than that of the other chains (50 Å^2^ vs 34 Å^2^); similarly, the average B-factor of V_H_ (chain H) in the tetragonal structure of the complex is higher than that of the other chains (79 Å^2^ vs 52 Å^2^). However, in both cases, the B-factors of the ZP2/IE-3 interface residues in the AHL and BXY complexes are comparable (30 Å^2^ vs 28 Å^2^ (crystal form I); 45 Å^2^ vs 42 Å^2^ (crystal form II)).

Structural alignments were performed using Chimera (19), Coot and PyMOL (Schrödinger, LLC). Protein-protein interfaces and oligomeric states were analyzed using, PIC (20) and PDBsum (21). All figures were created with PyMOL.

### Mouse gamete collection

3-to 5-week old C57BL6/J female mice were used as oocyte donors. Mice were kept under controlled light and temperature conditions with free access to food and water at the Preclinical Laboratory Portal South Core Facility of Karolinska University Hospital (Huddinge, Sweden). Ovarian stimulation was induced by intraperitoneal administration of 5 IU Pregnant Mare Serum Gonadotrophin (PMSG; Folligon, Intervet) and, 48 h later, 5 IU human Chorionic Gonadotropin (hCG; Chorulon, Intervet). 12 h after the hCG injection females were sacrificed and ampullas from individual animals were retrieved and equilibrated in HTF medium (Embryo Max, Millipore). Cauda epididymis were dissected from mature male mice and placed into 1 mL drops of HTF medium supplemented with 0.4% (w/v) BSA and covered with oil (OVOIL, Vitrolife). Sperm was capacitated by incubation for 1 h at 37ºC, 5% CO_2_. All animal experiments were performed in accordance with the approval of the local ethical committee (Stockholm South Animal Ethics Committee, Sweden; application number S-26-15).

### *In vitro* fertilization assays

IE-3 scFV and control hGH, purified in 10 mM Na-HEPES pH 7.5 and 150 mM NaCl, were concentrated to 1.2 mg mL^-1^ and 1.3 mg mL^-1^, respectively. After 15 min equilibration in HTF medium at RT, 5 μL protein was added to IVF drops containing an average of 16 oocytes surrounded by cumulus cells and covered with oil. Protein and oocytes were incubated for 1 h at 37ºC, 5% CO_2_. Because the fertilization block induced by IE-3 is concentration-dependent (22), a 10-fold excess of IE-3 scFV (corresponding to 2.2 μM final protein concentration in the IVF drop) was used to ensure full saturation of its binding sites in the ZP. Control hGH was used at the same molar concentration of IE-3 scFV. After capacitation, ∼5 × 10^6^ mL^-1^ highly motile sperm was added to the IVF drops containing oocytes and proteins; 4 h later, oocytes were washed, incubated overnight, and observed to assess 2-cell cleavage occurrence. Fertilization rates were calculated based on the number of healthy cells.

To count the number of sperm attached to the ZP, 5 groups of 5-10 oocytes (hGH) or 6 groups of 5-15 oocytes (IE-3 scFV), corresponding to a total of 36 or 54 oocytes, respectively, were washed in PBS 1 h after having added capacitated sperm and fixed in 2% (v/v) paraformaldehyde (PFA; Sigma-Aldrich). Oocytes were then washed three times and finally fixed using 2-cell embryos as washing controls. Sperm was counted by performing a through-focus series on each oocyte.

### Confocal microscopy

2 cell-embryos or unfertilized oocytes were fixed with 2% (v/v) PFA, washed with 100 μL PBS and blocked with a solution containing 1x PBS, 3% (w/v) BSA for 1 h at RT. After three washing steps with PBS, cells were incubated overnight at 4°C with anti-5His monoclonal (1:1000). The day after, after three washing steps in 1x PBS, cells were transferred into a drop containing PBS supplemented with 3% BSA and goat anti-mouse IgG Alexa 488 secondary antibody (1:1000; Thermo Fisher Scientific) and incubated for 1 h at RT. Samples were washed, transferred to a glass coverslip with PBS and imaging was performed on a Nikon A1R confocal microscope at RT, at the Karolinska Institutet Live Cell Imaging Core Facility (Huddinge, Sweden).

## Notes

### Competing Interest Statement

The authors have declared no competing interest.

## References

1. Bill & Melinda Gates Foundation. Accelerating Discovery for Non-Hormonal Contraceptives. (2020). Available at: https://gcgh.grandchallenges.org/challenge/accelerating-discovery-non-hormonal-contraceptives.

2. P. M. Wassarman, E. S. Litscher, Mouse zona pellucida proteins as receptors for binding of sperm to eggs. Trends Dev. Biol. 15, 1–13 (2022).

3. C. A. Shivers, A. B. Dudkiewicz, L. E. Franklin, E. N. Fussell, Inhibition of sperm-egg interaction by specific antibody. Science 178, 1211–1213 (1972).

4. J. F. Kirkpatrick, J. W. Turner Jr, I. K. Liu, R. Fayrer-Hosken, Applications of pig zona pellucida immunocontraception to wildlife fertility control. J. Reprod. Fertil. Suppl. 50, 183–189 (1996).

5. R. A. Fayrer-Hosken, D. Grobler, J. J. Van Altena, H. J. Bertschinger, J. F. Kirkpatrick, Immunocontraception of African elephants. Nature 407, 149 (2000).

6. W. Sun, Y. H. Lou, J. Dean, K. S. Tung, A contraceptive peptide vaccine targeting sulfated glycoprotein ZP2 of the mouse zona pellucida. Biol. Reprod. 60, 900–907 (1999).

7. M. A. Avella, B. Baibakov, J. Dean, A single domain of the ZP2 zona pellucida protein mediates gamete recognition in mice and humans. J. Cell Biol. 205, 801–809 (2014).

8. S. Nishio, et al., ZP2 cleavage blocks polyspermy by modulating the architecture of the egg coat. Cell 187, 1440–1459.e24 (2024).

9. I. J. East, D. R. Mattison, J. Dean, Monoclonal antibodies to the major protein of the murine zona pellucida: effects on fertilization and early development. Dev. Biol. 104, 49–56 (1984).

10. J. Li, et al., Vectored antibody gene delivery mediates long-term contraception. Curr. Biol. 25, R820–R822 (2015).

11. I. J. East, B. J. Gulyas, J. Dean, Monoclonal antibodies to the murine zona pellucida protein with sperm receptor activity: effects on fertilization and early development. Dev. Biol. 109, 268–273 (1985).

12. I. J. East, J. Dean, Monoclonal antibodies as probes of the distribution of ZP-2, the major sulfated glycoprotein of the murine zona pellucida. J. Cell Biol. 98, 795–800 (1984).

13. I. Raj, et al., Structural Basis of Egg Coat-Sperm Recognition at Fertilization. Cell 169, 1315–1326.e17 (2017).

14. E. Dioguardi et al., 9H4R: Molecular basis of immunocontraception: structure of mouse ZP2 ZP-N1 domain bound to fertilization-blocking monoclonal antibody IE-3 V_H_V_L_ (crystal form I). Protein Data Bank. https://www.rcsb.org/structure/unreleased/9H4R. Deposited 21 October 2024.

15. E. Dioguardi et al., 9H4R: Molecular basis of immunocontraception: structure of mouse ZP2 ZP-N1 domain bound to fertilization-blocking monoclonal antibody IE-3 V_H_V_L_ (crystal form II). Protein Data Bank. https://www.rcsb.org/structure/unreleased/9H4S. Deposited 21 October 2024.

## References

1. I. Raj, et al., Structural Basis of Egg Coat-Sperm Recognition at Fertilization. Cell 169, 1315–1326.e17 (2017).

2. M. Bokhove, et al., Easy mammalian expression and crystallography of maltose-binding protein-fused human proteins. J. Struct. Biol. 194, 1–7 (2016).

3. W. Sun, Y. H. Lou, J. Dean, K. S. Tung, A contraceptive peptide vaccine targeting sulfated glycoprotein ZP2 of the mouse zona pellucida. Biol. Reprod. 60, 900–907 (1999).

4. M. Monné, L. Han, T. Schwend, S. Burendahl, L. Jovine, Crystal structure of the ZP-N domain of ZP3 reveals the core fold of animal egg coats. Nature 456, 653–657 (2008).

5. I. J. East, J. Dean, Monoclonal antibodies as probes of the distribution of ZP-2, the major sulfated glycoprotein of the murine zona pellucida. J. Cell Biol. 98, 795–800 (1984).

6. J. Li, et al., Vectored antibody gene delivery mediates long-term contraception. Curr. Biol. 25, R820–2 (2015).

7. D. J. Leahy, C. E. Dann 3rd, P. Longo, B. Perman, K. X. Ramyar, A mammalian expression vector for expression and purification of secreted proteins for structural studies. Protein Expr. Purif. 20, 500–506 (2000).

8. D. de Sanctis, et al., ID29: a high-intensity highly automated ESRF beamline for macromolecular crystallography experiments exploiting anomalous scattering. J. Synchrotron Radiat. 19, 455–461 (2012).

9. P. D. Adams, et al., PHENIX: a comprehensive Python-based system for macromolecular structure solution. Acta Crystallogr. D Biol. Crystallogr. 66, 213–221 (2010).

10. W. Kabsch, XDS. Acta Crystallogr. D Biol. Crystallogr. 66, 125–132 (2010).

11. A. J. McCoy, et al., Phaser crystallographic software. J. Appl. Crystallogr. 40, 658–674 (2007).

12. V. Muñoz Robles, et al., Crystal structure of two anti-porphyrin antibodies with peroxidase activity. PLoS One 7, e51128 (2012).

13. A. Teplyakov, et al., On the domain pairing in chimeric antibodies. Mol. Immunol. 47, 2422–2426 (2010).

14. G. Bunkóczi, R. J. Read, Improvement of molecular-replacement models with Sculptor. Acta Crystallogr. D Biol. Crystallogr. 67, 303–312 (2011).

15. T. C. Terwilliger, et al., Iterative model building, structure refinement and density modification with the PHENIX AutoBuild wizard. Acta Crystallogr. D Biol. Crystallogr. 64, 61–69 (2008).

16. P. Emsley, B. Lohkamp, W. G. Scott, K. Cowtan, Features and development of Coot. Acta Crystallogr. D Biol. Crystallogr. 66, 486–501 (2010).

17. P. V. Afonine, et al., Towards automated crystallographic structure refinement with phenix.refine. Acta Crystallogr. D Biol. Crystallogr. 68, 352–367 (2012).

18. V. B. Chen, et al., MolProbity: all-atom structure validation for macromolecular crystallography. Acta Crystallogr. D Biol. Crystallogr. 66, 12–21 (2010).

19. E. F. Pettersen, T. D. Goddard, C. C. Huang, UCSF Chimera—a visualization system for exploratory research and analysis. Journal of (2004).

20. K. G. Tina, R. Bhadra, N. Srinivasan, PIC: Protein Interactions Calculator. Nucleic Acids Res. 35, W473–6 (2007).

21. R. A. Laskowski, et al., PDBsum: a Web-based database of summaries and analyses of all PDB structures. Trends Biochem. Sci. 22, 488–490 (1997).

22. I. J. East, D. R. Mattison, J. Dean, Monoclonal antibodies to the major protein of the murine zona pellucida: effects on fertilization and early development. Dev. Biol. 104, 49–56 (1984).

